# Reduced representation optical methylation mapping (R^2^OM^2^)

**DOI:** 10.1101/113522

**Authors:** Assaf Grunwald, Hila Sharim, Tslil Gabrieli, Yael Michaeli, Dmitry Torchinsky, Rani Arieli, Matyas Juhasz, Kathryn R Wagner, Jonathan Pevsner, Jeff Reifenberger, Alex R Hastie, Han Cao, Elmar Weinhold, Yuval Ebenstein

## Abstract

Reduced representation methylation analysis utilizes a subset of CpGs in order to report the overall methylation status of the probed genomic regions. Here, we use this concept in order to create fluorescent optical methylation profiles along chromosomal DNA molecules for epigenetic profiling. Reduced representation optical methylation mapping (R^2^OM^2^) in combination with Bionano Genomics next generation genome mapping (NGM) technology provides a hybrid genetic/epigenetic genome map of individual chromosome segments spanning hundreds of kilobase pairs (kbp). These long reads, along with the single-molecule resolution, allow for epigenetic variation calling and methylation analysis of large structural aberrations such as pathogenic macrosatellite arrays not accessible to single-cell next generation sequencing (NGS). We show that in addition to the inherent long-read benefits of R^2^OM^2^, it provides genomic methylation patterns comparable to whole genome bisulfite sequencing (WGBS) while retaining single-molecule information. The method is applied here to detect methylation along genes, around regulatory histone marks and to study facioscapulohumeral muscular dystrophy (FSHD), simultaneously recording the haplotype, copy number and methylation status of the disease-associated, highly repetitive locus on chromosome 4q.

## INTRODUCTION

DNA methylation, specifically methylation of the 5-carbon of cytosine, is the most studied and among the most significant epigenetic modifications^1^. In mammalian DNA, methylation mostly occurs on cytosines within CpG dinucleotides (DNA motifs where the cytosine is followed by a guanine residue) that are methylated to an extent of ~70%^2^. Approximately 60% of human gene promoters contain clusters of CpGs referred to as CpG islands (CGIs)^3^. CpG methylation plays an important role in regulation of gene expression with the general notion that hypermethyation of promoters suppresses gene expression^4^. Thus, the methylation status of gene promoters may predict gene activity and relate epigenetic transformations in development and disease to gene expression and protein abundance.

Another genetic feature regulated by CpG methylation is repetitive arrays. These variable copy number elements are homologous DNA sequences, exhibiting identity (or great similarity) to each other^5,6^. Many repetitive elements are mobile and can transpose across the genome, perform homologous recombination events, and promote dynamic genomic transformation^6–8^. This unstable nature explains their size variability, both among different individuals and between different cells of the same individual^6,9,10^. Typically arrays are characterized by the number of units composing them, which has been shown to affect their activity^11^. Methylation of repeat units adds another dimension of variability to these elements: Locally it may regulate the function of individual units and globally it can change the effective number of units in an array altering its activity. In this context, it has been shown that methylation levels of repetitive DNA can regulate repeat-related genetic diseases^12,13^, and are correlated with various types of cancer and their severity^14^. One striking example of a repeat array, addressed in this work, is facioscapulohumeral muscular dystrophy (FSHD), one of the most commons forms of muscular dystrophy, affecting approximately 1 in 7,000-20,000 individuals^15–18^.

The “gold-standard” method for studying CpG methylation is bisulfite sequencing, often used for whole-genome, base-pair resolution methylation profiling by next generation sequencing (WGBS). As such, it provides an averaged representation of the sample’s DNA sequence, with distinction between methylated and non-methylated cytosines^19^. The main shortcomings of bisulfite sequencing are the high cost and the requirement for large amounts of starting material due to chemical degradation of the DNA, and the need to split the sample for sequencing the treated and untreated samples separately.

Reduced representation bisulfite sequencing (RRBS) utilizes restriction enzymes that cut at CpG sites, thus specifically enriching for fragments that end within CpG islands. The fragment ends are ligated to adapters, size selected, bisulfite converted, and sequenced to give a representative genome-scale methylation profile. Using the restriction enzyme MspI to cut DNA at CCGG sites captures 60% of gene promoters, thus producing important regulatory information while requiring very little input sample^20^. The low input implies that fewer reads are required for accurate sequencing, allowing for high throughput, low-cost methylation analysis for clinical and single-cell applications^21,22^. Nevertheless, bisulfite sequencing and RRBS have limited accessibility to repetitive regions due to the inherent limitations of short-read sequencing, and are often unable to quantify the length and arrangements of the repeats ^23^ (Figure S2). Thus, in many cases the genetic and epigenetic characteristics of repetitive elements are still unknown. With this in mind, it is possible that some of the many NGS-inferred sequence duplications are in fact longer repetitive elements. Moreover, since both repetitive elements and methylation patterns tend to display somatic mosaicism, manifesting different structures in different cells^10^, their variability is often masked in the averaged NGS results.

Here we harness the reduced repre^10^sentation concept in order to access the methylation profile of long individual chromosome segments by optical genome mapping. Reduced representation optical methylation mapping (R^2^OM^2^) seeks to simultaneously capture large scale structural and copy number variants together with their associated methylation status.

DNA optical mapping^24–28^ stands out as an attractive approach for studying large genomic rearrangements such as repeat arrays. The approach consists of a set of techniques for stretching long genomic fragments, followed by imaging of these fragments using fluorescence microscopy. Image processing is used to read out a fluorescently labeled barcode along the molecules that provides genetic information such as the genomic locus of the imaged molecule as well as the size and number of large repeat units. The most advanced method for optical genome mapping involves linearizing and uniformly stretching of fluorescently barcoded DNA molecules in highly-parallel nanochannel arrays. This technique, commercialized by Bionano Genomics Inc., is capable of large scale genetic mapping and automated copy number analysis on a genome scale^29,30^.

The use of fluorescence microscopy presents the potential of obtaining several types of information simultaneously from the same DNA molecule by using different colors. Labeling different genomic features allows studying epigenetic marks and DNA damage lesions in their native genomic context on the single-molecule level^31–33^. In order to facilitate such multiplexing, we have developed an enzymatic labeling reaction that can distinguish methylated from non-methylated cytosines, and showed that it can be used to detect and quantify methylation levels in synthetic DNA molecules translocated through solid-state nanopores^34^. The bacterial methyltransferase M.TaqI in combination with a synthetic cofactor analogue can fluorescently label the adenine residue in TCGA sites containing non-methylated CpGs. M.TaqI has potential access to about 5.5% of all CpGs, however, these sites represent over 90% of gene promotors and hence M.TaqI is a good candidate for reduced representation methylation profiling and for studying genomic regulatory function. Combining this profile with a genetic pattern generated by labeling of specific sequence motifs, results in a hybrid genetic / epigenetic barcode for every DNA molecule. Using the genetic labels for mapping to the genome reference allows, for the first time, simultaneous assignment of the epigenetic labels to their specific genomic locations, and the study of DNA methylation patterns over large genomic fragments at single genome resolution.

We use R^2^OM^2^ to compare the methylation levels of primary white blood cells and a B-Lymphocyte derived cell line [NA12878], directly quantifying the reduction in CpG methylation levels after Epstein-Barr virus cell-line transformation. Furthermore, we show that optical methylation profiles are well correlated with whole genome bisulfite sequencing (WGBS) results and give good representation of the methylation status around genes and regulatory elements. In particular, optical signal levels from promotor methylation islands correlate with published gene expression data^35,36^. Finally, we demonstrate the utility of R^2^OM^2^ to simultaneously probe copy number and methylation level in macrosatellite arrays, that when contracted can result in disease, serving as a potential diagnostic assay for FSHD.

## RESULTS

### Methylation sensitive optical mapping quantifies genome-wide and locus specific methylation levels

The DNA methyltransferase M.TaqI methylates the adenine at TCGA sites. The enzyme may be “tricked” to incorporate a fluorophore instead of a methyl group by using a synthetic cofactor analogue^37–39^ (Figure S1a upper panel). However, the M.TaqI recognition site (TCGA) contains a nested CpG dinucleotide, which when methylated blocks the M.TaqI reaction^40^.

We first tested whether this methylation sensitive labeling reaction may be combined with optical mapping for studying the methylation levels of various genomes and specific DNA regions. We chose to compare primary blood cells to an Epstein-Barr virus (EBV) transformed blood lymphocyte cell-line, NA12878 (Coriell Institute, USA), where reduced genome-wide methylation levels have been reported^41^.

Genomic DNA was first nick-labeled with the nicking enzyme Nt.BspQI, to create a distinct genetic pattern of red labels along the DNA, regardless of the epigenetic state of the sample^42^. Next, a second layer of information was added by labeling only non-methylated CpGs with a green fluorophore using M.TaqI (Figure S1). The labeled DNA was loaded into nanochannel array chips, for dual-color optical genome mapping (Figure 1a). The genetic barcode (red labels) was used to align the molecules to their corresponding sequence locations. After alignment, the epigenetic labels were used to generate R^2^OM^2^ profiles, displaying levels of non-methylated CpG sites along the human genome, with resolution of 1kbp (Figure 1c). We used the IrysView software suite (Bionano Genomics Inc.) for automatic label detection and genomic alignment of the imaged molecules. Our goal was to provide global quantification of the relative methylation levels in the two samples. As expected^41^, EBV immortalized cell line genomes were hypomethylated compared to DNA from primary cells with a 3.6 fold higher number of M.TaqI labels compared to the primary DNA (16.0 Vs. 4.7 labels per 100 kbp; Figure 1b). These results demonstrate the utility of M.TaqI labeling for quantitative assessment of methylation levels in various samples.

**Figure 1.**
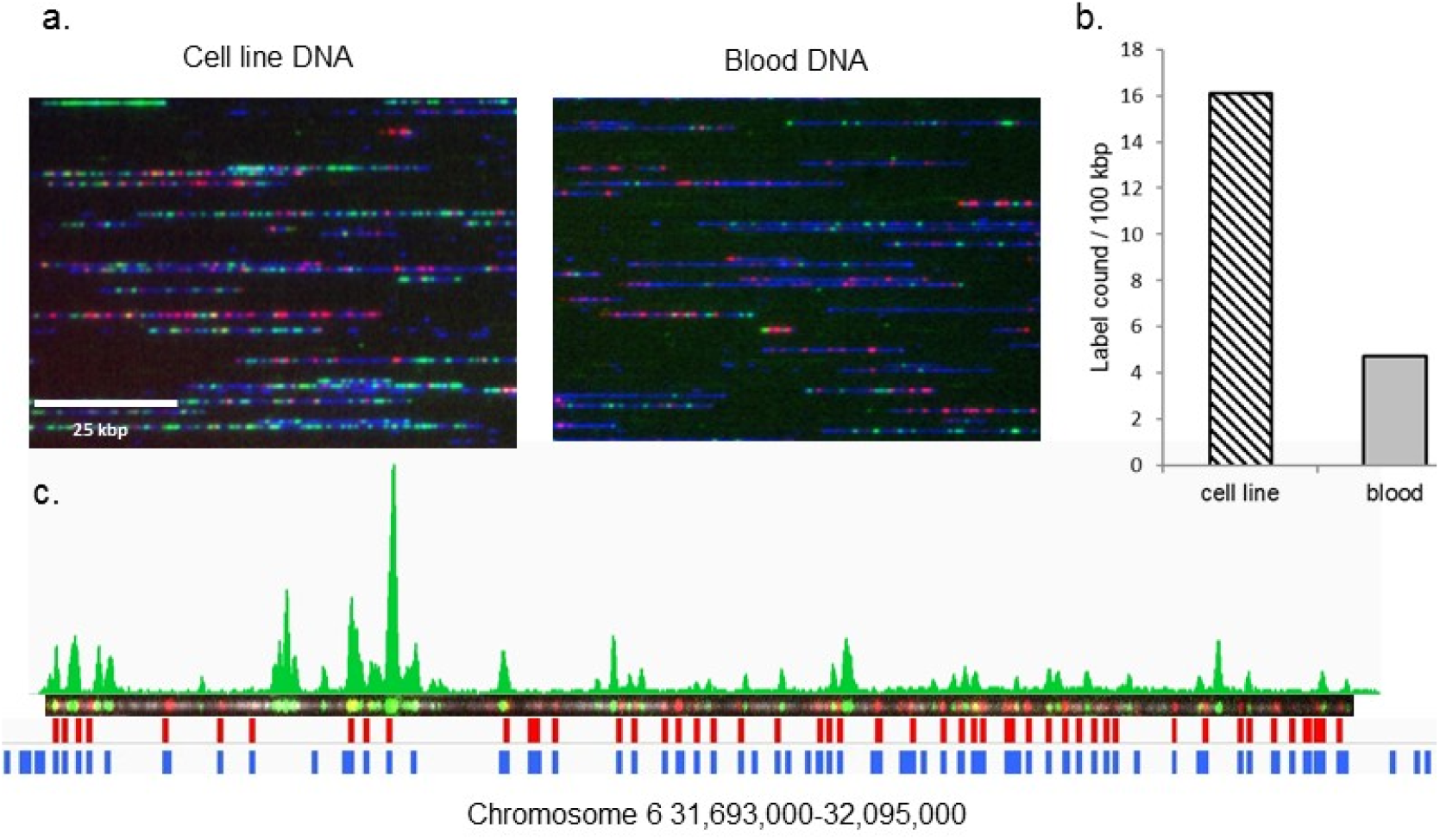
Methylation detection on genomic DNA. a. Genomic DNA from human lymphocyte cell line culture (left image) or primary human blood cells (right image) was dual labeled with genetic labels (Nt.BspQI, red labels) and methylation sensitive labels (M.TaqI, green labels), and imaged on a nanochannel array chip. Representative images from both samples are presented here. DNA backbone, stained with YOYO-1 is displayed in blue. **b.** Non-methylated CpG labels were detected for both DNA samples. The bar-graph presented here displays number of detected labels per 100 kbp (in total 4.6 Gbp were sampled for the primary blood cells DNA and 1.2 Gbp for the lymphocyte cell line). Dashed bar displays label number for the cell line DNA and gray bar for the primary blood DNA. It is clear that epigenetic labels are more frequent in the immortalized cell-line genome, indicating that it is relatively non-methylated as expected. **c.** A representative dual labeled molecule from the primary blood sample. R^2^OM^2^ profile along the molecule is presented in green above the molecule image. The detected locations of genetic labels are shown in red below the image, and the reference barcode, used for alignment to the locus in chromosome 6 is shown in blue.

### R^2^OM^2^ provides genome-wide methylome profiles of genes and regulatory elements

We validated the quality of R^2^OM^2^ profiles by correlating our results to available “gold standard” WGBS data. Levels of non-methylated CpGs were calculated along gene bodies and promotor regions and around four commonly used regulatory histone marks (Figure 2). In all cases the results showed high correlation with WGBS data and is in agreement with the expected methylation status of these regions^43^. Additionally, we found that on a genome scale, R^2^OM^2^ profiles correlate well with WGBS profiles, especially given the inherent 5.5% sampling rate (Figure 2c).

**Figure 2.**
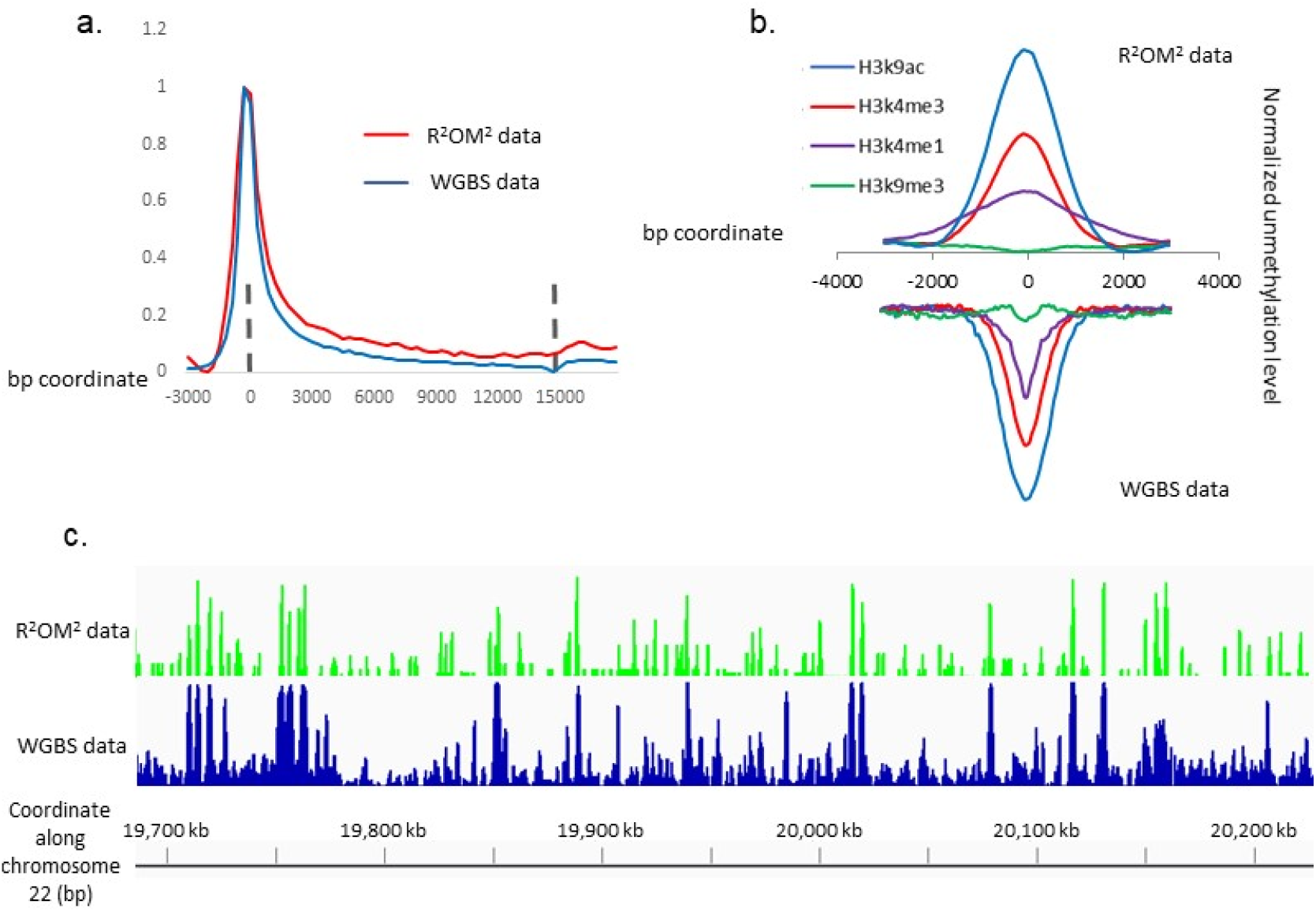
R^2^OM profiles correlate with WGBS based profiles. **a.** Non-methylated CpG levels along gene bodies for R^2^OM^2^ data (red) and WGBS data (blue). For both data types, non-methylated CpG levels were sampled along each gene separately and the plot displays average and normalized non-methylated CpG values for all genes in the human genome. The presented region spans gene body and 3 kbp up and downstream from the transcription start and end sites (indicated by dashed lines at the left and right rims of the track respectively). It is clear that both methods produce highly similar tracks reporting depleted methylation around the transcription start sites as expected. **b.** Plots of methylation levels spanning 6 kbp around four regulatory histone marks. For each histone modification, the genome-wide average methylation values are plotted for each modification type: H3K4me3 (red), H3K9me3 (green), H3K4me3 (purple), H3K9ac (blue). For the sake of demonstration, the WGBS plots are inverted so that they appear below the R^2^OM^2^ data. It is clear that both data types produce similar results not only in the shape of non-methylation profiles around the regulatory sites but also in their intensities. **c.** Non-methylated CpG levels as detected along the human genome: R^2^OM^2^ (green track) and WGBS (blue track). A region of about 500 kbp is presented to allow sufficient resolution (the presented region was randomly chosen along chromosome 22 and reflects the overall correlation).

The main advantage of R^2^OM^2^ is the ability to generate extremely long reads from individual molecules, thus, allowing for direct comparison between long methylation patterns ranging hundreds of kbps. This is exemplified in Figure 3 showing R^2^OM^2^ profiles from genomes of three primary white blood cells aligned to the same 250 Kbp genomic region in chromosome 1p. The R^2^OM^2^ profiles are presented below the calculated distribution of CpG and M.TaqI sites in this region and the positions of associated genes are presented in the bottom panel. The molecules in Figure 3d exhibit similar methylation profiles that are well correlated with positions of calculated CpG islands. Small variations in the pattern are seen and can be attributed to common variation in methylation status among white blood cells^44^. Furthermore, the maps allow relating promoter methylation to gene expression data. For instance, a CpG island that lies in the promotor region of the PHF13 gene (indicated by a red box), was found to be non-methylated by both sequencing and R^2^OM^2^, and is known to be expressed in white blood cells^36^. In contrast, the promoter of the *KLHL*^41^ gene (indicated by a black box), was detected as methylated by both methods. This muscle specific gene is indeed expected to be silenced in blood cells^35^. These findings and their agreement with established NGS results validate the ability of R^2^OM^2^ to unravel the methylome at a single-cell-like level. Optical mapping allows genomic information to be read directly from individual, un-amplified, long fragments of DNA, thus mitigating the bias of large structural and copy number variation in single cell sequencing experiments.

**Figure 3.**
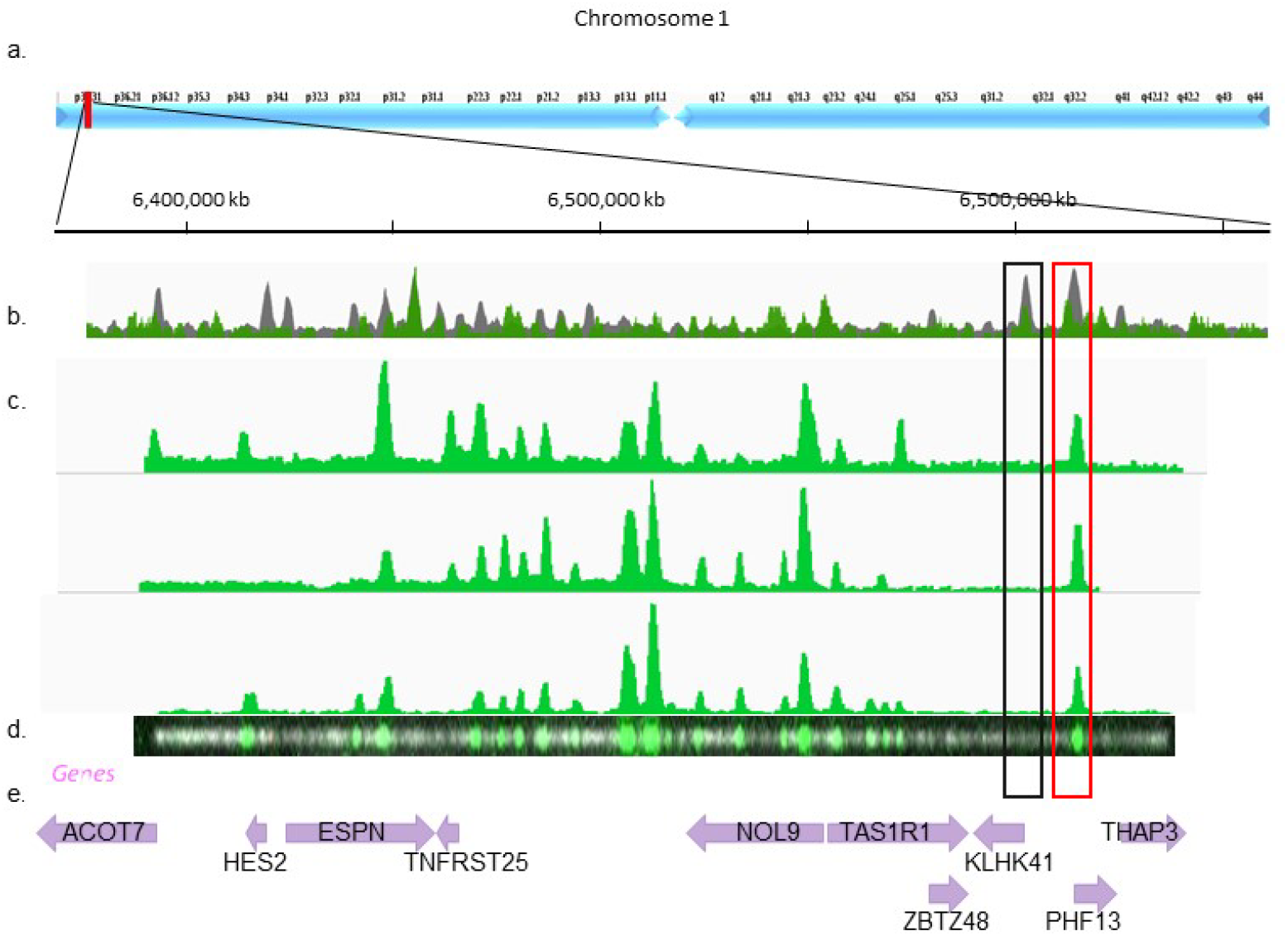
**a.** Detailed view of chromosome 1:6,350,000-6,670,000 bp. **b.** A histogram of CpG (gray) and M.TaqI (green) sites across the region. **c.** Digital representation of three detected molecules aligned to the reference sequence based on Nt.BspQI labels (P-value<10^−20^). Label intensity is shown in green. **e.** The raw image of the backbone (light gray) and the M.TaqI channel (green) of the bottom molecule presented in d. **f.** Gene body locations and corresponding HGNC gene symbols. Each gene is represented as a purple arrow and the direction of the arrow indicates gene orientation. Black and red rectangles across b-e indicate methylated and non-methylated gene promoters, respectively.

### Simultaneous quantification of copy number and methylation state in DNA tandem repeats

Macrosatellite arrays, repetitive DNA that spans up to millions of base pairs across the genome, are extremely challenging for analysis by next-generation sequencing. Analysis is further complicated by the recent understanding that DNA methylation plays a crucial role in the function of these regions. One striking example of the significance of methylation in such regions is the D4Z4 array on chromosome 4, which is directly related to the muscular dystrophy FSHD^45^. Recent evidence show that both the number of D4Z4 repeats and their methylation status constitute the genotype of the disease, dictating whether the individual manifests disease symptoms or not. Commonly, healthy individuals carry an array of more than 10 repeats. However, even long arrays result in FSHD symptoms when losing their methylation (termed FSHD2) while carriers of short but highly methylated repeat arrays, do not manifest the disease^15,16^. The multiple combinations of copy number and methylation level span a broad range of possible variations which are correlated with the disease severity and manifestation ^16,17^.

To demonstrate the capabilities of optical mapping for simultaneous copy number quantification and DNA methylation detection, we studied a model system of FSHD-associated D4Z4 repeats from healthy human individual cloned into the CH16-291A23 bacterial artificial chromosome (BAC) (Figure 4a). We first attempted to evaluate the copy number of the studied array by NGS read-depth analysis^46^. We reasoned that the ratio between number of reads representing the repeat unit, relative to the number of reads detected for a single copy region, would provide the array copy number. Purified BAC DNA was sequenced using Illumina MiSeq to a read depth of 15,000X. We used read depth analysis to assess the D4Z4 copy number. Briefly, all sequencing reads were aligned to the BAC's reference sequence, containing only one repeat, and the copy number was calculated as the ratio between the number of reads (“coverage”) aligned to the repetitive and the non-repetitive regions respectively. The coverage along the non-repetitive sequence displayed variation of up to 25%, while the coverage along the repeat sequence was extremely variable, averaging 63% of the mean read depth (Figure S2). The ratio between the median coverage value along the repetitive and non-repetitive sequences was ~8, implying that this is the number of D4Z4 repeats along the BAC (Figure 3c). However, the large standard deviation values strongly demonstrate the unreliability of this method, as well as the sensitivity of PCR amplification and NGS to the exact content of the investigated sequence.

**Figure 4.**
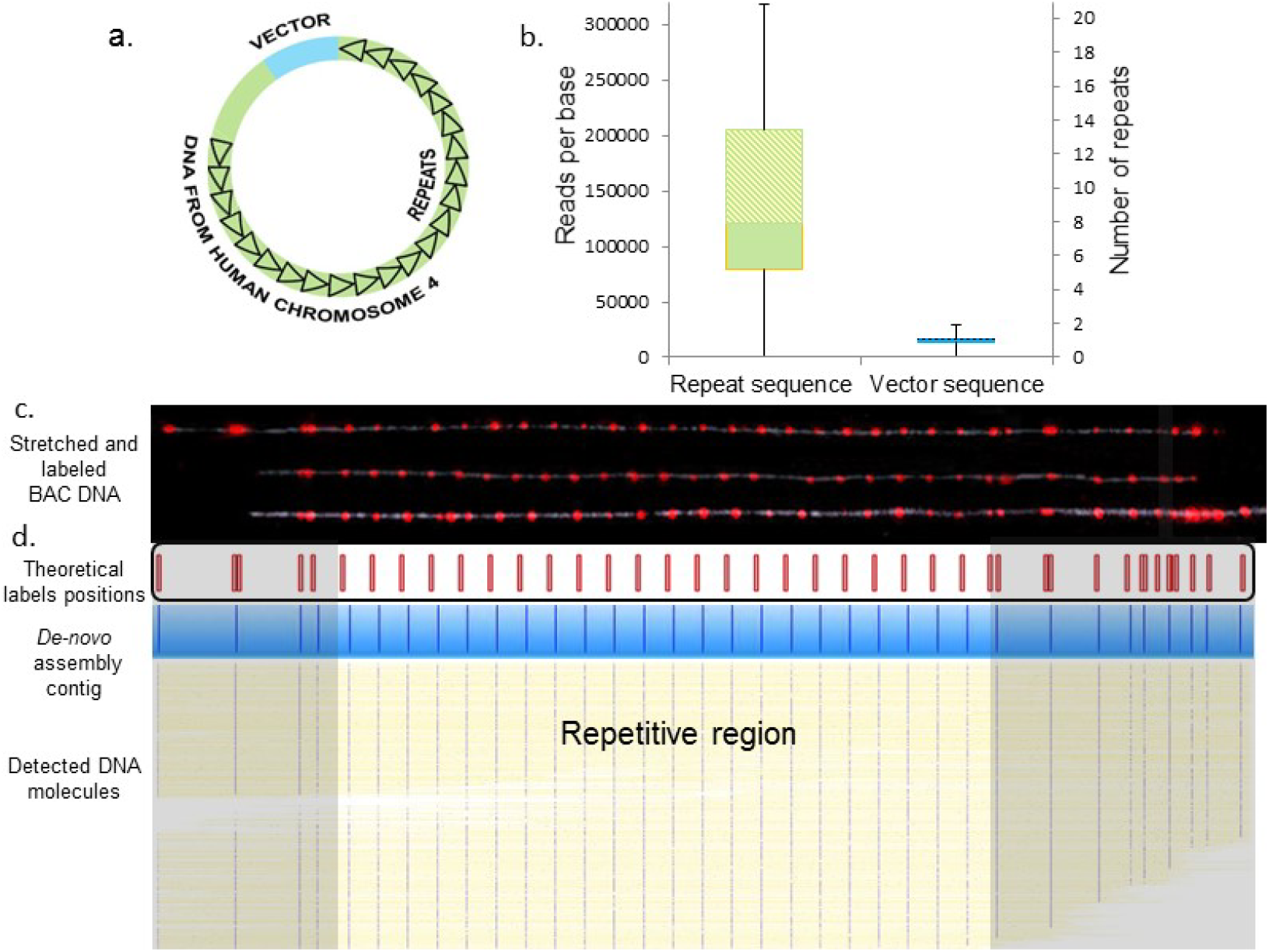
Copy number analysis. **a.** The FSHD BAC model system contained an unknown number of D4Z4 repeats (black triangles), an unknown size of genomic DNA 5' to the repeat array (green), and the cloning vector (blue). **b.** Read depth analysis box plots displaying the 25th percentile (bottom of the lower box), the median (intermediate between boxes) and the 75th percentile (top of the upper box), for coverage values of the sequencing reads along the repeat region (left box), and the non-repetitive region (including the vector and the upstream to repeat region, right box). The scale on the right is normalized to the median coverage along the non-repetitive region. **c.** Representative images of three intact model molecules labeled with Nb.BsmI (red dots) and stretched on modified glass surfaces. The pattern of labeled molecules can be aligned to the reference map presented below the images (expected labeling locations are shown in red). The repetitive region can be distinguished by virtue of the equally spaced labels, each representing a repeat, and quantified by simply counting the labels (23 repeats). **d.** 627 digital representations of labeled DNA molecules stretched and imaged in a nanochannel array chip. The detected molecules were *de-novo* assembled into a single consensus map (displayed in blue) which shows excellent agreement with the theoretical map. The yellow lines represent the DNA backbone and the blue dots represent the detected labels. The assembled map contained 23 repeat units.

Next, we directly visualized the repeats along stretched DNA molecules. We used tailored labeling of sequence motifs contained in the repetitive element to highlight individual repeats and allow for physical counting of the copy number. We nick-labeled the BAC DNA using nicking enzymes (Nb.BsmI or Nb.BssSI) which have a single recognition site on the 3.3 kbp long D4Z4 repeat sequence, yielding a single distinct fluorescent tag for each repeat unit. The labeled DNA was stretched and immobilized for visualization on modified glass slides, using a simple micro-fluidic scheme (see supporting information). This method allowed fluorescence imaging of the entire DNA contour and localization of individual fluorescent labels along the DNA (Figure 4c). The same sample was loaded onto an Irys instrument (Bionano Genomics Inc., CA. USA), which facilitates high-throughput DNA stretching and imaging in nanochannel array chips. The post-imaging analysis, performed by the IrysView software suite, involved automatic label detection and *de-novo* assembly of the molecules into a contiguous consensus barcode. The resulting consensus map was created in an unsupervised manner based on label patterns from approximately one thousand detected molecules^47,48^. When comparing the non-repetitive region of this contig to the one predicted from the known sequence, an almost perfect match was obtained (p-value <10^−43^, Figure 4d). The consensus repetitive region is unambiguously composed of 23 D4Z4 repeats, demonstrating the potential of the technique for genetic diagnosis of FSHD.

For methylation analysis, we performed R^2^OM^2^ as an overlay on the repetitive genetic barcode. We used red fluorophores for the genetic barcode and green fluorophores for methylation mapping. Figure 5a shows the unique pattern created by M.TaqI along the non-methylated BAC^49^, highlighting non-methylated repeat units. M.TaqI has two close-by recognition sites on each repeat unit that result in one visual label on each repeat, due to diffraction limits. Nevertheless, when over-stretching the non-methylated BAC on modified microscope slides, the two green methylation labels flanking the red genetic label were clearly resolved, in agreement with the theoretical dual-color barcode (Figure 5b).

**Figure 5.**
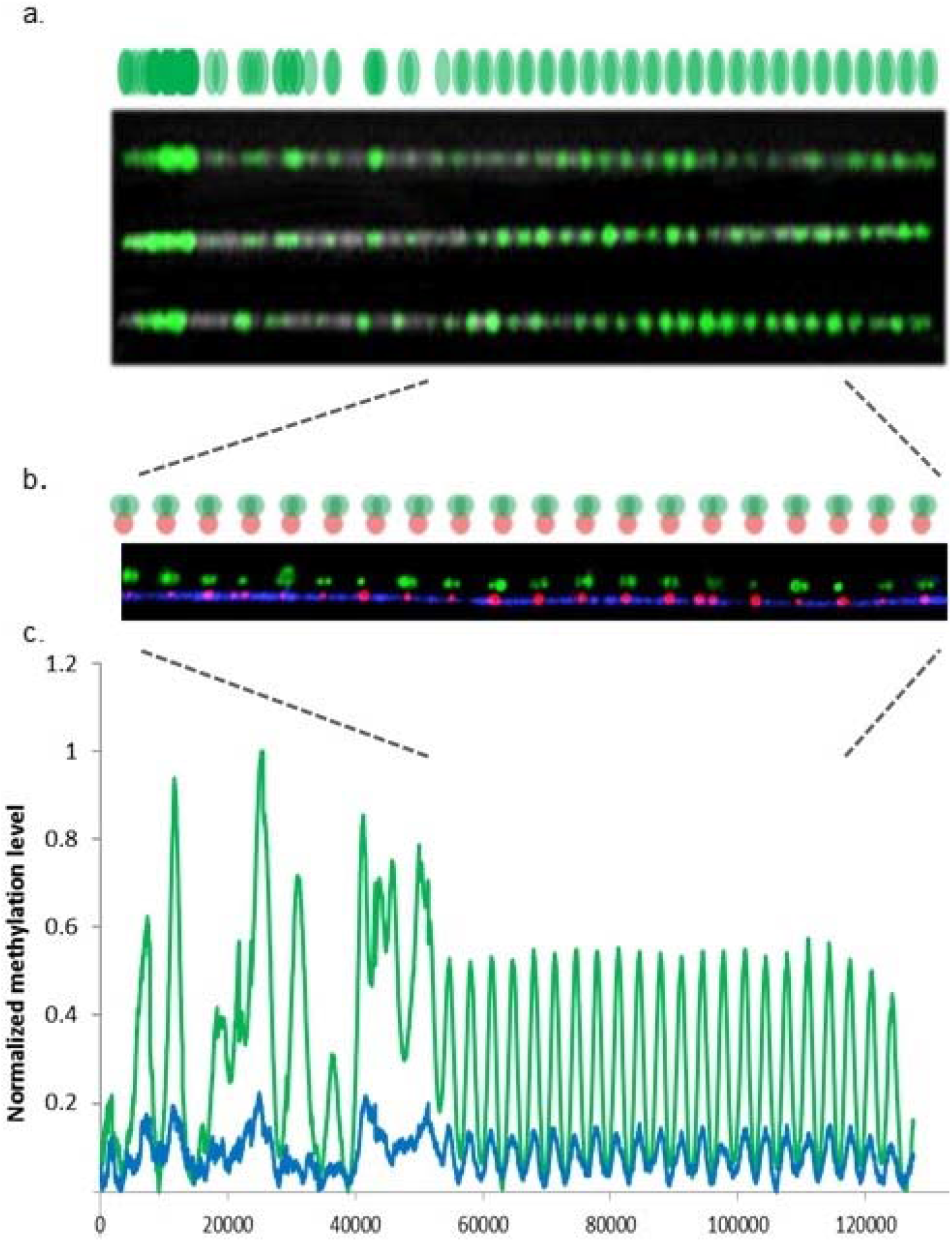
**a.** A reference map simulating the relative expected locations of R^2^OM^2^ labels generated by M.TaqI along the stretched BAC (green). Below, images of three non-methylated, M.TaqI labeled BAC molecules linearized and stretched in nanochannels and aligned to the reference map. **b.** Partial genetic map (red) and R^2^OM^2^ (green) from the repetitive region of the FSHD BAC stretched on a modified glass slide. The genetic identity and the number of repeat units are highlighted by labeling with the nicking enzyme Nb.BsmI (red labels). Co-labeling the DNA with M.TaqI (green labels) indicated that the molecules were non-methylated. The displayed image is an overlay of red and green channels along the repetitive region of a single BAC molecule. Above the repeats is the reference map for the region. The green channel is shifted upwards to allow better visualization. The higher stretching factor achieved on modified glass compared to nanochannels, enables detection of the two M.TaqI labels flanking the genetic label in each repeat. **c.** Comparative R^2^OM^2^ of non-methylated and partially-methylated BAC samples. Normalized integrated maps of detected M.TaqI labels are presented for the non-methylated (green, 18074 molecules) and partially-methylated (blue, 9089 molecules). Both plots highly correlate with the expected reference map.

To simulate the native state of DNA, where repeat arrays are methylated to variable degrees, we partially methylated the DNA using the CpG-specific DNA MTase M.ssSI. We repeated the dual-labeling reaction on the partially methylated sample, as well as on a non-methylated control, and analyzed both on the nanochannel array chips. After image analysis, we used the red genetic labels for automated *de-novo* assembly and generated the consensus map with the 23 repeats as described earlier. With thousands of molecules now aligned to the consensus map, we could compare the green methylation patterns generated by R^2^OM^2^ on the non-methylated and partially methylated samples. Figure 5c shows integrated methylation profiles generated from the molecules detected in each data set. The non-methylated sample shows frequent labeling, at the expected locations (green dots), while the partially methylated DNA sample shows significantly fewer labels (blue dots). It is clear from the integrated plot that in the partly methylated sample, methylated CpGs are distributed equally among the repeat units, in line with the fact that the partial methylation was random and equal for all repeats in the array. These results clearly demonstrate that R^2^OM^2^ enables not only single-molecule and single-repeat resolution but also assessment of the average methylation status for each individual repeat in the array across a population of different DNA molecules.

### R^2^OM^2^ enables genetic / epigenetic diagnosis of FSHD

Finally, we performed R^2^OM^2^ on whole blood DNA extracted from a donor previously diagnosed with FSHD via commercial testing based on pulsed-field gel electrophoresis and Southern blot. Figure 6 shows the genome reference barcode (green bar) and the two haplotypes detected in the sample (blue bars). Genetic labeling allowed distinguishing between the two very similar 4qA and 4qB alleles, each with a distinct copy number of D4Z4 repeats. Macrosatellite repeats were visually marked by the equally spaced methylation sensitive labels (marked green) indicating a non-methylated array. We measured 4 repeat blocks for the 4qA allele and 8 repeat blocks on the non-pathogenic 4qB allele^50^. This was similar to previous genetic assessments based on clinically performed pulsed-field gel electrophoresis in which the EcoRI/BlnI allele of 15 kb on chromosome 4qA corresponding to approximately 4 repeats and the 4qB allele was estimated to have a EcoR1/Blnl allele size of 27 kb, corresponding to approximately 7 D4Z4 repeats. These data demonstrate the utility of this optical mapping approach for characterizing the genetic and epigenetic profile of FSHD as well as other macrosatellite arrays and large structural variants.

**Figure 6.**
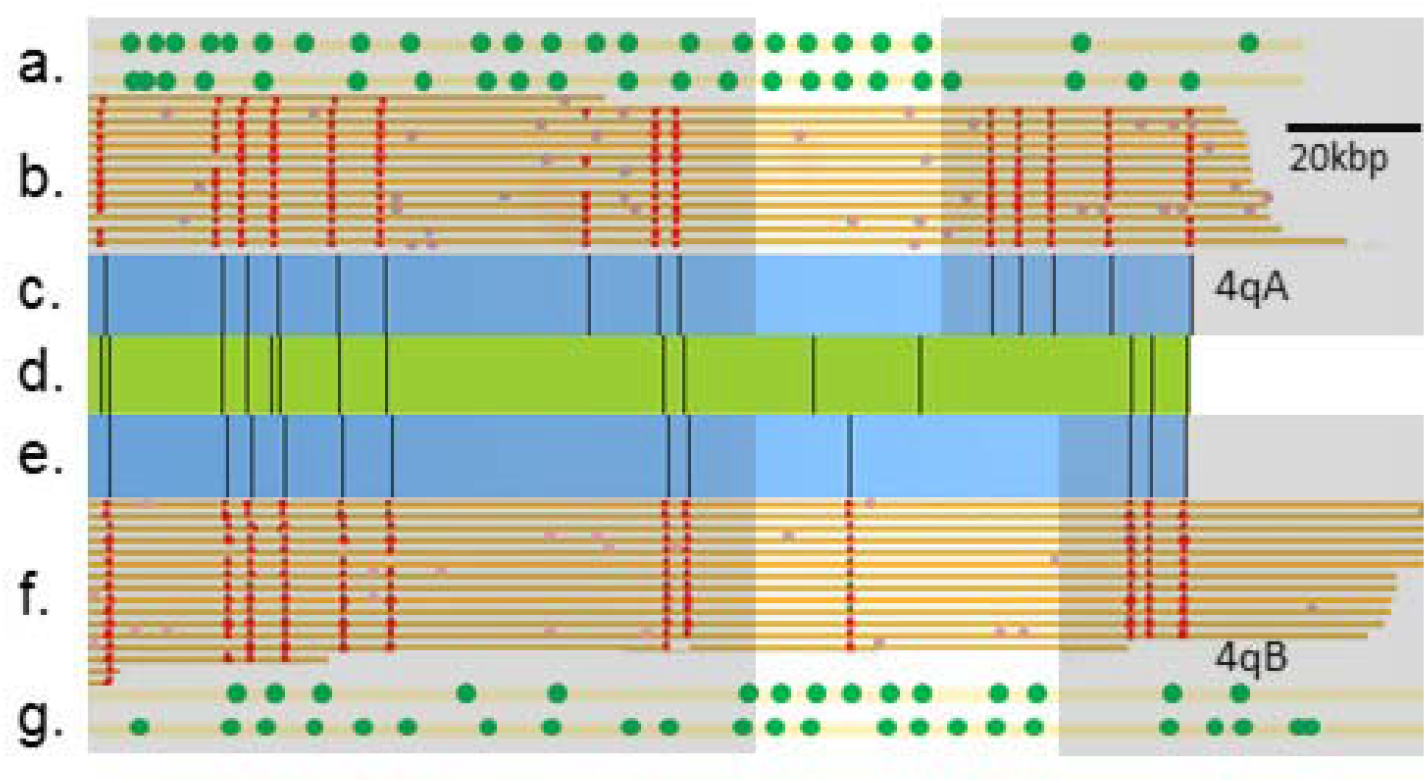
Structural variation analysis and methylation analysis of the pathogenic contraction on chromosome 4q of a patient with FSHD. The central green horizontal bar **(d.)**, shows the reference map for chromosome 4q. The consensus optical maps of the two haplotypes, 4qA **(c.)** and 4qB **(e.)**, generated *by De-Novo* assembly are above and below the reference. **b./f.** show genetic maps generated from individual molecules aligned to the genome reference. Maps were clustered by haplotype (4qA and 4qB). Red dots represent labels used for alignment. **a./g.** show R^2^OM^2^ profiles of DNA molecules aligned to the 4qA/4qB allele and displaying hypomethylation of the D4Z4 repeats. The gray background covers the non-repetitive region on each allele.

## DISCUSSION AND OUTLOOK

Optical genome mapping has recently been used to map whole human genomes at high coverage and to highlight genetic variability between individuals with unprecedented detail^29,30^. This work adds an epigenetic component to optical maps, providing, for the first time, correlated genetic/epigenetic profiles for individual DNA molecules spanning hundreds of thousands of base pairs. Utilizing state-of-the-art optical genome mapping technology, combined with DNA MTase-assisted methylation detection, we create a reduced representation optical methylation map, R^2^OM^2^, which reports on the methylation status of 90% of gene promoters. The reduced representation provides sufficiently sparse labeling to comply with the low resolution of optical maps while providing high correlation with BS-seq data around regulatory elements. We demonstrate automatic, high-throughput analysis of global methylation levels in various cell types by quantifying the amount of non-methylated CpGs. Additionally, R^2^OM^2^ provides long read / low-resolution methylation profiles at different genomic loci with emphasis on CpG islands and regulatory sequence elements such as gene promoters and histone binding sites. This property may allow studying the epigenetic profiles of genes together with remote regulators such as distant enhancers and cis elements. One area in which R^2^OM^2^ has a particular advantage over existing technologies is the study of DNA methylation of large structural variants and repetitive arrays. DNA repeats are dynamic regions, exhibiting high variability both in length and methylation status, and thus may differ significantly between individuals^9,51^. Today, in the era of personalized medicine, it is becoming increasingly accepted that the full profile of genomic structural variation, including DNA repeats, is directly linked to health and susceptibility to disease^8,52^. R^2^OM^2^ offers single-molecule level information on the size of the region, the number of repeat units, and the methylation status of individual repeats. This detailed information is inaccessible via current genome technologies such as NGS, DNA arrays, and quantitative PCR, which mostly provide averaged or inferred data and cannot specifically address individual repeat units.

We demonstrate the utility of R^2^OM^2^ for characterizing the D4Z4 repeat array, where both the size and the methylation status of the repeats affect FSHD disease manifestation^16^. Specifically, the reported approach can distinguish between healthy individuals, FSHD1, and FSHD2 carriers by combining copy number and methylation level information. Moreover, the detailed view of the methylation status of individual repeats may offer new insights into the mechanism of disease and may lead to a more individualized prognosis than is offered by current commercial testing. Notably, the R^2^OM^2^ concept is not limited to D4Z4 repeats, and various labeling enzymes may be used to address different target sequences, thereby extending the repertoire of available targets for this method. Further development of this technique may serve to map differentially methylated single-molecule patterns on a genome-wide scale, potentially allowing simultaneous genetic/epigenetic haplotyping, as well as ultra-sensitive detection of epigenetic transformations.

## ONLINE METHODS

### Human Subjects

This study was approved by The Johns Hopkins School of Medicine Institutional Review Board. The donor with a clinical diagnosis of FSHD1 was confirmed by the University of Iowa Diagnostic Laboratories to have a contracted D4Z4 array on a 4qA allele by pulse-field gel electrophoresis and Southern blotting. The EcoRI/BlnI allele of 15 kb on chromosome 4qA corresponds to approximately 4 repeats, and of 27 kb allele on chromosome 4qB corresponds to approximately 7 repeats.

### DNA samples

λ-Bacteriophage DNA (New England BioLabs, Ipswich MA, USA) was used as provided. BAC DNA was purified from *E.coli* cells containing the CH16-291A23 BAC. Cells were cultured overnight in LB containing 12.5 µg/ml cloramphenicol (Sigma-Aldrich, Rehovot, Israel) at 30°C. BAC DNA was purified from the cells using the NucleoBondXtra BAC kit (MACHEREY-NAGEL Inc. Düren, Germany). DNA from a lymphocyte cell line (NA12878, Coriell Institute for Medical Research, Camden New Jersey, USA), from primary blood cells (purchased from HemaCare Inc. Van Nuys, CA, USA) and whole blood DNA from human subjects were purified in agarose plugs in order to maintain large DNA fragments, following the Bionano Genomics IrysPrep protocol^29^.

### M.TaqI - assisted labeling for R^2^OM^2^

To generate methylation sensitive labels we used the DNA MTase M.TaqI, which catalyzes the transfer of a carboxytetramethylrhodamine (TAMRA) fluorophore from the synthetic cofactor AdoYnTAMRA onto the adenine residue within its recognition sequence (TCGA)^53^. The labeling reaction was carried out as follows: 1 µg of DNA was reacted with 37.5 ng of M.TaqI and 40 µM of AdoYnTAMRA in labeling buffer (20 mM Tris-HOAc, 10 mM Mg(Cl)2, 50 mM KOAc, 1 mM DTT, pH 7.9) in the presence of 0.01% Triton-X 100 and 0.1 mg/ml BSA in a total reaction volume of 25 µL at 60°C for 1 hour. The labeled DNA was reacted with 40 µg of proteinase K (Sigma-Aldrich, Rehovot, Israel) at 45°C for 1 hour to disassemble protein-DNA aggregates. For methylated samples CpG methylation was performed prior to labeling using the CpG-specific DNA MTase M.SssI (Thermo Scientific, Waltham, MA, USA) according to manufacturer’s instructions but with twice the suggested amount of enzyme to ensure complete methylation. To obtain partial methylation, the reaction was carried out using the recommended amount of enzyme, but for 75% of the recommended incubation time. Methylation was verified by digestion with a methylation-sensitive restriction enzyme HpaII (New England BioLabs Inc., Ipswich MA, USA), followed by gel electrophoresis to ensure that the DNA was protected or partially protected from restriction (Figure S1).

### Nick labeling for optical genome mapping

DNA was prepared in a nick-labeling-repair reaction (NLR) involves (a) the nicking enzyme Nb.BsmI or Nt.BspQI which generates single-strand nicks at its specific recognition sites (GAATGC or GCTCTTCN respectively), (b) a DNA polymerase enzyme incorporates fluorescent nucleotides at the nicked sites; and finally, (c) a DNA ligase enzyme repairs the remaining single-strand breaks. For the NLR reaction involving Nt.BspQI, DNA was labeled using the IrysPrep kit (Bionano Genomics Inc., San Diego CA, USA) according to manufacturer’s instructions. For Nb.BsmI-based NLR, DNA (900 ng) was first reacted with 30 units of the enzyme (New England BioLabs Inc., Ipswich MA, USA) in 30 µl NEBuffer 3.1 for 120 minutes at 65°C. Next, the DNA was reacted with 15 units of Taq DNA polymerase (New England BioLabs Inc., Ipswich MA, USA) in the presence of the following nucleotides: dGTP, dCTP dATP (Sigma-Aldrich, Rehovot, Israel) and the fluorescent nucleotide dUTP-Atto647 (Jena Bioscience GmbH, Jena, Germany) at a final concentration of 600 nM (each). The reaction was carried out in a reaction buffer (ThermoPol buffer, New England BioLabs Inc., Ipswich MA, USA) in a total volume of 45 µl for 60 min at 72°C. Finally, the DNA was reacted with 120 units of Taq DNA ligase (New England BioLabs Inc., Ipswich MA, USA) with 0.5 mM NAD^+^ (New England BioLabs Inc., Ipswich MA, USA), in a reaction buffer (ThermoPol Buffer, New England BioLabs Inc., Ipswich MA, USA) including 0.5 mM NAD+ (New England BioLabs Inc., Ipswich MA, USA) and of 10 µM dNTP mix, in a total reaction volume of 60 µL for 30 min at 45°C. For R^2^OM^2^ experiments, DNA was initially labeled by NLR and then 0.05 µg - 0.5 µg of the labeled DNA was reacted with M.TaqI as described above (the reaction was scaled down accordingly). Prior to M.TaqI labeling the Nb.BsmI NLR reaction products were re-embeded in 1% agarose plug to allow it’s washing in water (plugs were incubated for ten minutes in water, this was repeated three times by replacing the water). In the case of Nt.BspQI-NLR, the M.taqI reaction was performed in the NLR buffer with addition of 1x Buffer 4 and 1x BSA (New England BioLabs Inc., Ipswich MA, USA) and Ph was adjusted to 8.3 using 0.1M NaOH. Before imaging the agarose matrix was digested using GELase enzyme (Epicenter, Madison, WI, USA).

### Sample preparation, DNA stretching and imaging

Post labeling, BAC and λ DNA were cleaned by ethanol precipitation as has been described previously^53^. Genomic DNA was cleaned by embedding it into agarose plugs and washing these in TE buffer (see supporting information). Prior to imaging, the labeled DNA was stained with 0.5 µM of YOYO-1 (Invitrogen, Carlsbad, CA, USA) for visualization of its contour. DTT (Sigma-Aldrich, Rehovot, Israel) was added to the reaction (200 mM) to prevent photo bleaching and DNA breaks. To stretch the DNA from its random coil conformation into a linear form, allowing imaging of its contour, we used two types of experimental scheme: The first type of stretch used modified glass surfaces. In this approach, DNA sample-solutions were flowed by applying capillary forces or using microfluidics, on glass surfaces that were chemically modified to facilitate the DNA's anchoring and stretching on the surfaces (see supporting information). After stretching, the DNA was imaged using an epifluorescence microscope (FEI Munich GmbH, Germany) equipped with a high-resolution EMCCD IXon888 camera (Andor Technology Ltd. Belfast, UK). A 150 W Xenon lamp (FEI Munich GmbH, Germany) was used for excitation with filter sets of 485/20ex and 525/30em, 560/25ex and 607/36em, and 650/13ex and 684/24em (Semrock Inc., Rochester NY, USA) for the YOYO-1, TAMRA, and Atto-647 channels respectively. For high-throughput mapping experiments, samples were analyzed in nanochannels array chips (Bionano Genomics Inc., San Diego CA, USA). On the chip, DNA is forced into 45 nm square nanochannels using an electric field and is stretched along the channel axis for imaging. This process is carried out in automated cycles by the Irys instrument (Bionano Genomics Inc., San Diego CA, USA) ^42^.

### Data analysis

For analysis of the high-throughput nanochannel array data, raw images were processed and DNA molecules were detected and digitized by custom image-processing and analysis software (Irys Extract^54^ or IrysView^29^). Genetic labels were assigned one set of coordinates along the molecules (the genetic map) and the methylation labels are assigned another set (the methylation map). One output of the detection process is the number of labels per 100 kbp allowing for direct comparison between labeling levels of different samples, and quantitative assessment of methylation levels (Figure S3).

For R^2^OM^2^, DNA molecules are first mapped to the genome reference based on the match between the fluorescent genetic pattern along the molecule and the pattern expected from the known reference sequence. after alignment, the methylation maps indicated the distribution of non-methylated CpGs along the mapped genomic regions, and methylation profiles were exported in BED format for visualization in standard genome browsers such as the UCSC genome browser ^55^ (https://genome.ucsc.edu/index.html).

Details on bioinformatics analysis and comparison to BS sequencing data is provided in the supporting information

IrysView also enabled *de-novo* assembly of the detected molecules into a consensus barcode or contig (genome map) and its comparison with the theoretical barcode. This unsupervised assembly was used to assess the number of repeats in the FSHD BAC model system.

Images of DNA molecules stretched on modified glass surfaces were manually aligned, according to the barcode created outside the repetitive region. This enabled detection of the starting point of the repeat array, allowing counting of individual repeat units.

### Next-generation sequencing

Purified BAC DNA was sheared using Covaris AFA (Covaris Inc. Woburn MA, USA) and the fragments were size-separated by electrophoresis using agarose gel, allowing selective extraction of fragments within the range of 150-300 bp. Illumina sequencing libraries were prepared using NEXTFlex kit (Bioo Scientific Corporation, Austin TX, USA) and sequenced using MiSeq (Illumina Inc. San Diego, CA. USA) to a paired-end coverage of 15,000X. Sequencing reads were *de-novo* assembled using CLC Workbench software (CLC Bio-Qiagen, Aarhus, Denmark).

## ACKNOWLEDGEMENTS

We thank Prof. D. Gabellini for his kind gift of the CH16-291A23 BAC and K. Glensk for preparing M.TaqI. We acknowledge financial support from the German-Israeli foundation [I-1196-195.9/2012] (E.W. and Y.E.), the BeyondSeq consortium [EC program 63489] (Y.E.), the European Research Councils starter grant [337830] (Y.E.) i-Core program of the Israel Science Foundation [1902/12] (Y.E.), NIH U54HD060848 (KRW), and NIH U01 MH106884 (JP).

